# The influence of learning on robustness of working memory to distractors

**DOI:** 10.1101/2022.03.08.482959

**Authors:** Ermolova Maria, Beliaeva Valeriia, Novikov Nikita, Fedele Tommaso, Moiseeva Viktoria, Gutkin Boris, Feurra Matteo

## Abstract

Inhibition of irrelevant information is a crucial process within the working memory (WM) system. In the present study, we explored how inhibition of distractors is learned within a Sternberg-like WM task and how this learning process affects readout performance. We introduced distractors of varying strength as well as novel stimuli to a classic Sternberg task. Our findings show that the length of the training period affected obstructiveness of distractors as well as differences between WM loads. Significant differences between conditions observed early on in accuracy became void after general ceiling performance was reached. Moreover, accuracy and reaction times, which are commonly used as readouts of WM performance, were affected differently by training. While both improved significantly, only accuracy exhibited differential dependency of experimental conditions on training. It is essential to consider these dynamics when introducing a WM task which requires a long training period.

## 1. Introduction

Numerous models and theories have been established identifying building blocks that constitute working memory (WM). One commonality of these theories is the substantial role of attentional processes (Baddeley, 2012, D’Esposito & Postle, 2015, Eriksson et al., 2015). Attention is employed during the early perceptual stage of encoding, as well as during later stages of information processing (Awh et al., 2006). Efficiency of attentional selectivity is crucial for encoding, maintenance, and manipulation of information (D’Esposito & Postle, 2015).

One of the key features of selective attention is the protection of memory from distractors. WM has been shown to have a limited capacity for the amount of information being stored (Baddley, 2012, Luck & Vogel, 1997). The higher the load of information is, the lower is the task performance that requires its manipulation (Oberauer, 2001, Bays & Husain, 2008). Therefore, it is essential to control and filter what enters the memory and what, on the other hand, is ignored as noise. If a task involves a distractor, it places a higher demand on WM resources, possibly due to their overloading (Zanto & Gazzaley, 2009). Indeed, failure to ignore unnecessary information results in decreased WM performance (Zanto & Gazzaley, 2009, Vogel et al., 2005). Presence of a strong distractor during memory retention periods results in worse performance as compared to the case when a distractor is easier to ignore (Bonnefond & Jensen, 2012). Moreover, the processes of competition for a spotlight can influence the distractor’s disruptiveness in terms of allocation of attention (Oh & Kim, 2003). If a behaviorally relevant item is present in memory, it guides the attention, and attenuates the distractor’s effect. If, on the other hand, the item is too ambiguous to guide the attention, the distractor wins the competition. In line with the aforementioned studies, Fockert et al. (2001) showed that memory load is correlated with the ability to inhibit distractors: greater memory load leads to failed suppression of distractors. Nevertheless, such ability is also influenced by individual differences in the capacity to selectively encode information. Participants with high capacity are more efficient in filtering relevant items from irrelevant ones, while low capacity individuals encode more information, including both relevant items and distractors (Vogel et al., 2005).

The ability to inhibit distracting information during WM can be studied, among other ways, using the Sternberg WM paradigm (Sternberg, 1966). Participants are typically presented with a set of items to remember. Then, upon being presented with a probe, they have to identify whether the probe was part of a previously shown set. Distracting stimuli may be introduced during the task periods when items have to be retained in memory. This modification of the paradigm allows studying behavioral and neurophysiological mechanisms of inhibitory control during WM (Jensen et al., 2002).

While various studies implementing the Sternberg task use different types of stimuli (letters, objects, characteristics such as color and direction, etc.), the choice of the stimuli may influence the results of the experiment. In Bonnefond & Jensen (2012), the consonant letters of the English alphabet were used for memory and strong distractor stimuli, while punctuation marks were employed as weak distractors. The authors found that strong distractors led to a decrease in performance as reflected by longer response times compared to weak distractors. Moreover, cortical oscillations over parieto-occipital regions in the alpha band increased in power and phase-adjustment prior to the appearance of the distractor. This increase was higher for the condition that required inhibition of strong distractors. These results can be explained by the fact that strong distractors were perceptively more similar to the stimuli that had to be memorized, thereby more difficult to inhibit with respect to weak distractors. However, another property of stimuli that has to be considered is their semantic saliency. Indeed, letters of the participants’ native language vocabulary were employed as stimuli, which can be easily recognized and encoded, as well as be susceptible to chunking. It was shown that letters work as automatic priming while being presented for as brief as 20 ms (Jacobs & Grainger, 1991).

Here we investigate the ability of humans to ignore unfamiliar distracting information while being engaged in a behavioral WM task. Distractors of different complexity were introduced: stimuli that are harder to ignore due to perceptual similarity with the to-be-memorized items (e.g. same category) and the ones that are easier to ignore due to perceptual differences with respect to the to-be-memorized items (e.g. different category). Importantly, a newly developed set of stimuli which are novel and designed to be void of semantic meaning to participants was adopted. In particular, the influence of learning on WM performance was investigated.

We ran two behavioral experiments as follows. Experiment 1 was conducted in a single day. Participants were instructed to memorize a set of 4 figures, ignore the 5^th^ figure (i.e. a distractor) and match the 6^th^ figure (i.e. a probe) to the memorized set. Distractors fell into two categories: visually similar to memorized stimuli, and visually different from them. In a control condition no distractor was presented and participants were instructed to memorize a set of 5 figures and match the 6^th^ figure to the memorized set. According to our hypothesis: 1) the condition with a distractor that is visually similar to memorized stimuli should be harder to complete than the condition with visually different distractors; 2) the condition with the greatest number of elements for memorization should be the hardest to complete. Both conditions with distractors were expected to be easier than the condition with no distractor but a bigger memorization set, if the distractors were correctly ignored (as the memory load for the latter is larger).

The ability to inhibit distractors may be subject to the learning process. In Experiment 2, we explored the ability to ignore irrelevant information as a function of training. For that, a 2-day experiment was conducted with the same protocol as Experiment 1. We tested the impact of training, done on two consecutive days, on the ability of distractor inhibition during the WM task.

## 2. Experiment 1

### 2.1 Methods

#### 2.1.1 Participants

The data was collected from 30 right-handed participants 18 to 32 years old (22 females, mean age +/- SD 23.4 +/- 3.9). 5 participants were excluded due to technical reasons (e.g. incorrect understanding of the task, software malfunction), leaving N = 25 for the analysis. All participants had (1) no neurologic or psychiatric medical history; (2) normal (or corrected to normal) vision; (3) sufficient sleeping time the night prior to an experiment (as self-reported); (4) no intake of psychoactive substances the day before an experiment. The study was approved by the ethics committee of the Higher School of Economics, all participants provided written informed consent and received monetary compensation after the experiment.

#### 2.1.2 Sensitivity analysis

Sensitivity analysis was performed with G*Power assuming a one-way repeated-measures ANOVA on the effect of the distractor condition. It indicated a minimum effect size of eta_p_^2^ = 0.12 that can be reliably estimated with the obtained sample size (80% power, alpha = 0.05, nonsphericity correction = 1, correlation among repeated measures assumed to be 0).

#### 2.1.3 Stimuli and apparatus

Stimuli for memorization were abstract figures, unfamiliar to participants. Stimuli were created from several ancient alphabets (Fig. 1), modified and adjusted in size, line thickness and occupied space. These stimuli also served as strong distractors (i.e. distractors which are relatively hard to ignore).

**Figure 1.**
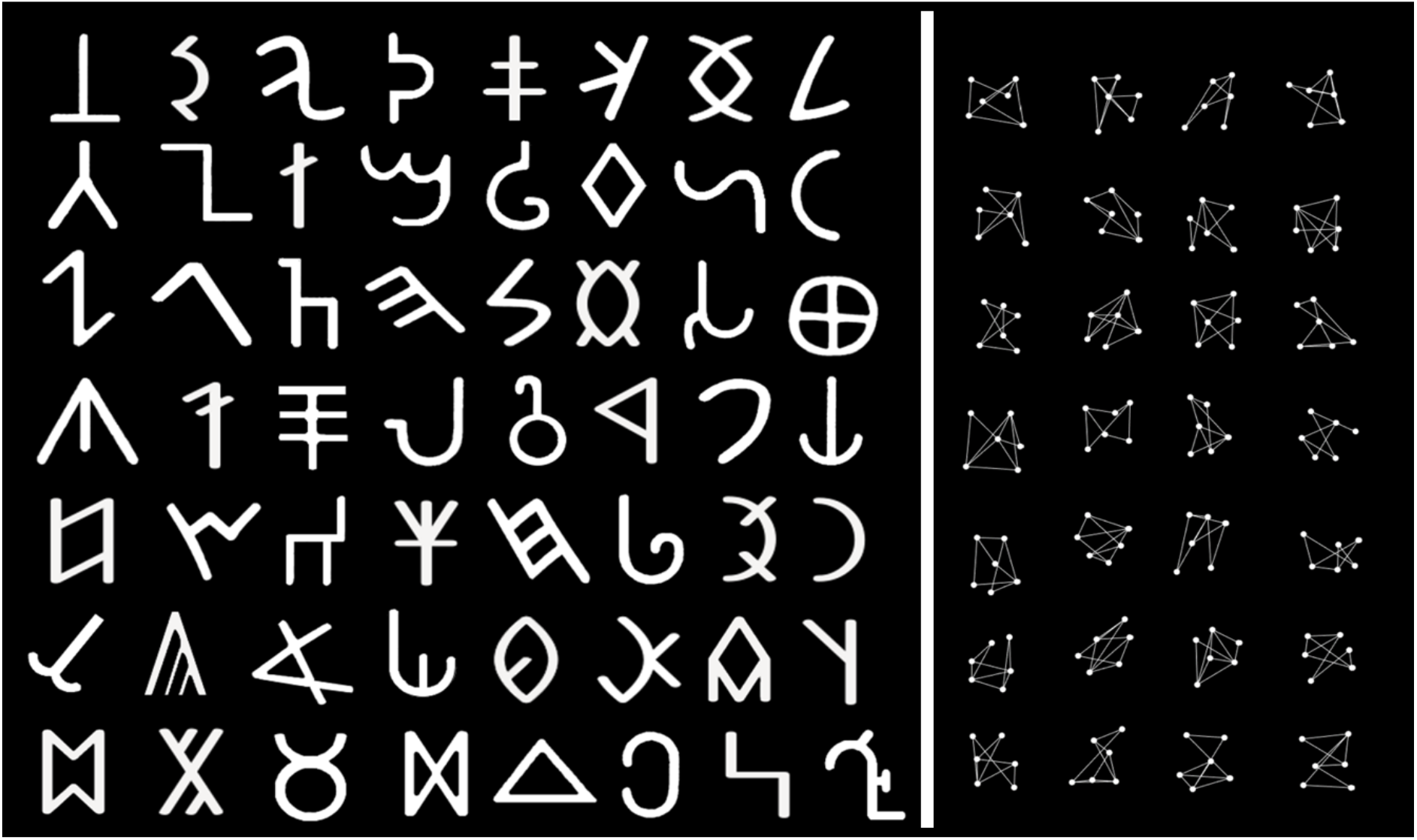
Stimuli and strong distractors - figures from ancient Brahmi, Phoenician, Orkhon alphabets, modified and adjusted to each other. Graphs with unilaterally connected 6 nodes served as weak distractors.

Weak distractors (i.e. easy-to-ignore distractors) were images of unilaterally connected 6-node graphs (Fig. 1). The final pool of stimuli consisted of 56 abstract figures and 28 graphs.

Stimuli were presented in white on black background on a screen positioned ~60 cm from the participant (1920×1080 screen resolution, 60 Hz refresh rate). Screen luminosity was reduced to 50% to preserve the eyes from fatigue. The experiment was built and run within the E-Prime 2.0 Professional program, presentation time lags were tested and deemed negligible (<0.015 sec).

#### 2.1.4 Task and procedure

The experimental paradigm of the study was a modified version of the Sternberg task (Fig. 2A). Six items were presented sequentially at the center of the screen for 0.033-sec each, with 1.1-sec intervals. First four items were part of a memorization set, while the fifth item was either part of the memorization set or a distractor, depending on the condition. The sixth item was a probe, upon demonstration of which the participant had to reply whether it was a member of the memorized set or not. The probe stayed on the screen until the participant gave an answer by pressing on one of the two buttons (labeled “Old” and “New”). Accuracy and reaction times were recorded after each answer as behavioral measures of success. In the strong and weak distractor conditions, the fifth item was a distractor to be ignored, visually similar or dissimilar to the memory set respectively. In the control condition (load-5), the fifth item was part of a memorization set and no distractor was presented.

**Figure 2.**
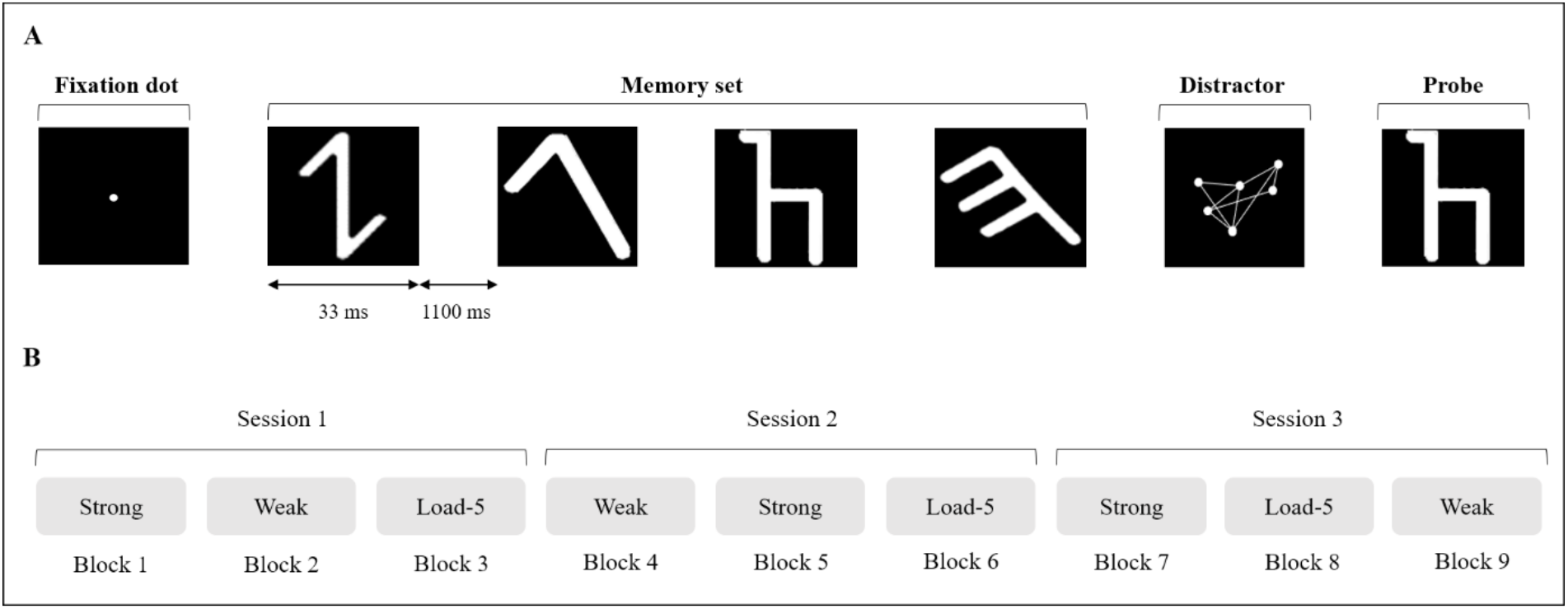
Experimental paradigm. A. Depiction of a trial. B. Experimental procedure for Experiments 1 and 2. Example from 1 participant.

Every trial started with a fixation dot at the center of the screen, staying there for a random period between 0.5 and 1.5 sec.

The experiment was divided into three sessions, each session including all three experimental conditions in a counterbalanced block design, with nine blocks in total (Fig. 2B). The nine experimental blocks consisted of 60 trials each (adding up to 180 trials per experimental condition and 540 trials overall). Averaged accuracy and reaction time were presented three times per each block (i.e. every 20 trials). The stimuli were randomized within trials, blocks, and sessions. Controlled randomization was performed using Matlab R2015a.

The appearance of all stimuli and distractors in trials was constrained in the following ways. The probe was matching one of the stimuli in the set in 50% of trials. Maximum sequence of the same consecutive trials (matching or non-matching) was constrained. The frequency of appearance of every stimulus within each block followed an approximately uniform distribution. Stimuli were never repeated within one memory set. Stimuli were never repeated in the same position within trials within each block. Distractor never matched stimuli of the memory set or the probe within the same trial.

An experiment began with a 10-trial training for each of the three experimental conditions. Both hands of the participant were always on the keyboard over the buttons labeled “Old” and “New” (positioned over “K” and “D” keys on a standard US keyboard layout). Positions of the answer buttons in respect to each other (i.e. right - left) were counterbalanced between participants.

An experiment ended with an open-answer questionnaire about memorization strategies and changes in them (see Supplementary).

#### 2.1.5 Data Analysis

The effects of weak-, strong-distractor and 5-load no-distractor condition on both accuracy (i.e. proportion of right answers) and response times (RTs) were tested. RTs were analysed only for successful trials (i.e. hits and true rejections). To avoid recency effect, both accuracies and RTs for the load-5 condition were calculated excluding trials in which the fifth item matched the probe. A two-way repeated measures ANOVA with a three-level factor Condition (Strong, Weak, Load-5) and a three-level factor Session (1, 2, 3) was used for statistical analysis. The threshold for statistical significance was set to p = 0.05. Greenhouse-Geisser correction was applied to the data which violated the assumption of sphericity. Post-hoc pairwise comparisons were performed via Student’s t-tests for the factors that reached significance. Bonferroni-Holm correction was applied to the post-hoc results to control for multiple comparisons. The analysis was conducted using R 4.1.0 (R Core Team, 2021) and statistical package “rstatix” (Kassambara, 2021).

### 2.2 Results

The goal of Experiment 1 was to verify the effects of distractors of varying complexity on WM as well as to evaluate the performance of Sternberg paradigm with regard to newly created stimuli.

There was no significant interaction between the effects of Session and Condition on accuracy (F(4,96) = 1.22, p = .31, eta_p_^2^ = .05) or RTs (F(4,96) = 1.04, p = .39, eta_p_^2^ = .04).

Condition had a significant effect on both accuracy (F(2,48) = 4.8, p = .013, eta_p_^2^ = .17) and RTs (F(2,48) = 4.63, p = .014, eta_p_^2^ = .16) (Fig. 3). Pairwise comparisons showed significant difference in accuracy between Weak and Load-5 conditions (t = 3.07, p < .01) but not between Strong and Weak or Strong and Load-5 conditions. In case of RTs, Weak condition was associated with significantly shorter RTs compared to Strong (t = 2.66, p = .01) and Load-5 conditions (t = 3.07, p < .01).

**Figure 3.**
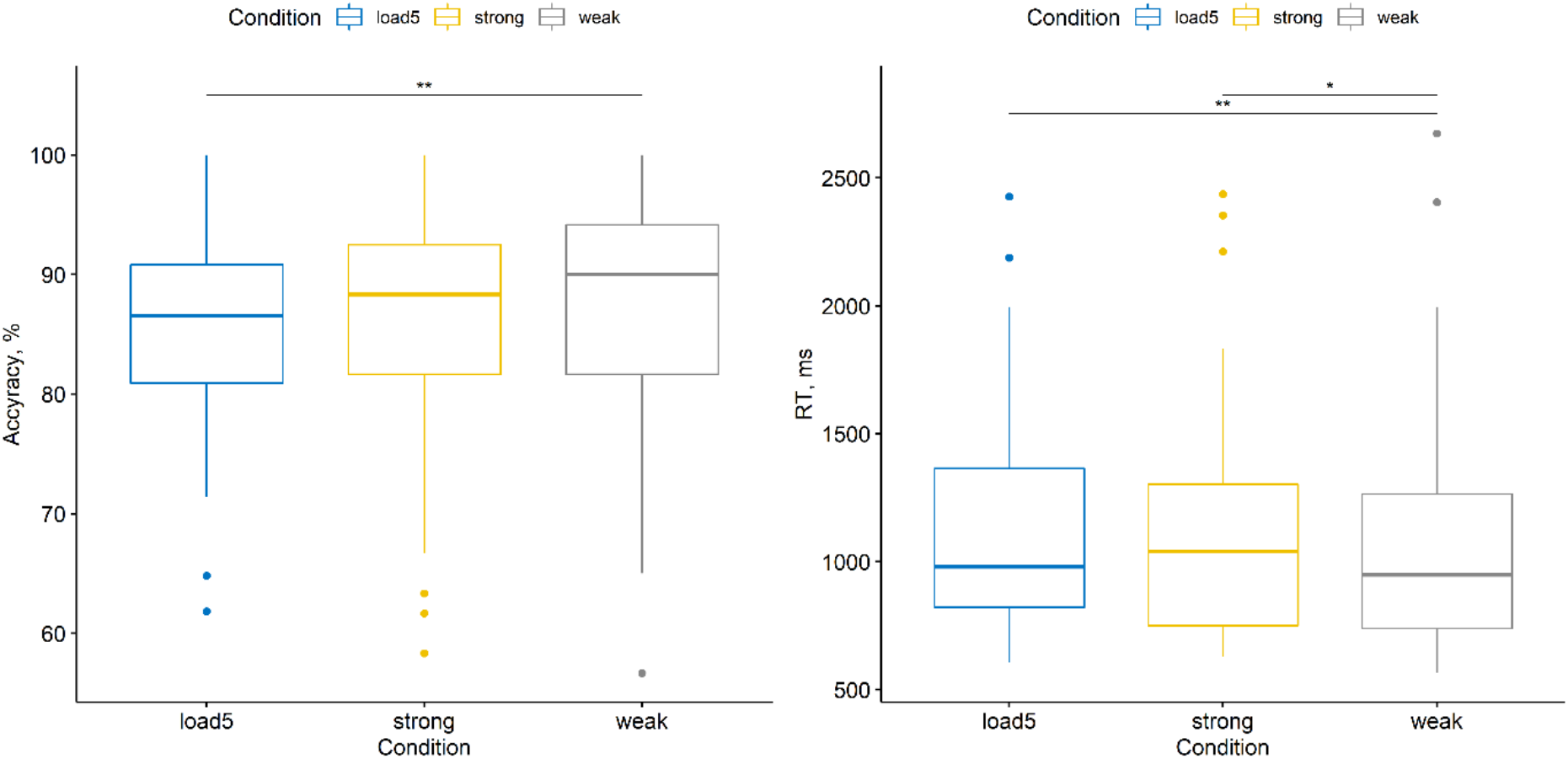
Experiment 1: Differences between distractor conditions in accuracy and RTs. *p<.05,**p<.01.

Session number had significant influence on both accuracy F(2,48) = 12.53, p < .001, eta_p_^2^ = .34) and RTs (F(2,48) = 26.36, p < .001, eta_p_^2^ = .52) (Fig. 4). For accuracy, the effect was prominent for the first session compared to the remaining sessions (t(1-2) = −4.62, p < .001; t(1-3) = −4.76, p < .001) but subsided for the two remaining sessions. RTs, on the other hand, exhibited a significant decrease throughout the whole experiment (t(1-2) = 5.94, p < .001; t(1-3) = 8.34, p < .001; t(2-3) = 4.32, p < .001).

**Figure 4.**
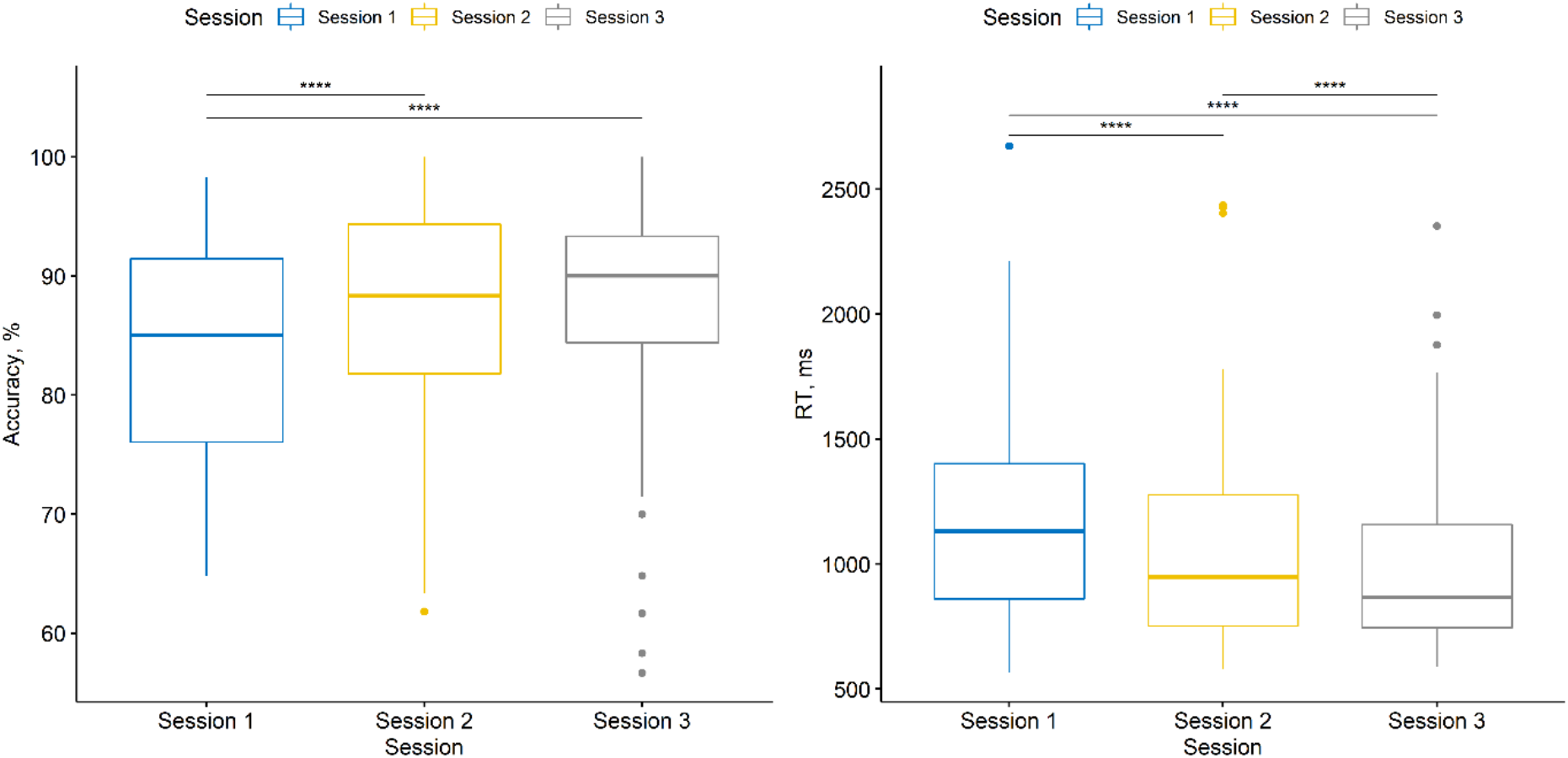
Experiment 1: Differences between experimental sessions in accuracy and mean RTs. ****p<.0001.

The post-experimental questionnaire allowed us to estimate memorization strategies of the participants and, in particular, changes in these strategies. Two strategies came forward: semantic (64%) and perceptual (4%) memorization. Semantic memory was based on making associations between the figures and familiar objects - participants reported naming a figure by a name of an object which it resembles and continuously repeating the names until the appearance of the probe (Craik & Lockhart, 1972). This pool of participants also reported ignoring the distractor by not naming it. Participants categorized under the “perceptual memorization” group memorized the image (i.e. the visual form) of the figure and evaluated whether the image of the probe looked similar to any figure of the set. 32% of participants reported changing the strategy during the experiment or using a mixed strategy. As the questionnaire was free-form, further exploration of the effect of memorization strategies on performance was deemed unsuitable.

### 2.3 Summary

Both accuracy and RTs were significantly modulated by distractor conditions. However, upon pairwise comparisons, significant differences between conditions with strong and weak distractors were observed in RTs but not in accuracy. Additionally, no significant differences were found between strong distractor 4-load and no-distractor 5-load condition in either accuracy or RTs. This result indicates that either the subjects failed to perform the task correctly (i.e. did not ignore the strong distractor) or that ignoring the strong distractor was as hard as memorizing an additional item. Significant influence of the session number on both accuracy and RTs indicated presence of a training effect. While accuracy remained stable after the first session, RTs were affected by training throughout all three sessions. For this reason, we conducted Experiment 2 which was a 2-day version of Experiment 1. The second day was introduced to give participants sufficient time to reach their ceiling performance in order to investigate possible interference of the training effect with the effects of main experimental conditions.

## 3. Experiment 2

### 3.1 Methods

#### 3.1.1 Participants

For Experiment 2, the data was collected from 18 participants from 19 to 29 years old (11 females and 7 males, mean age +/- SD = 23.7+/-3.3).

#### 3.1.2 Power analysis

Power analysis was performed assuming a one-way repeated-measures ANOVA on the effect of the distractor condition based on the effect sizes achieved in Experiment 1. It indicated that with a sample size of Experiment 2 the effect of the distractors on accuracy and RT will be detected with power of 0.82 and 0.79, respectively (alpha = 0.05, nonsphericity correction = 1, correlation among repeated measures assumed to be 0). The analysis was performed post-hoc after completion of Experiment 2.

#### 3.1.3 Task and procedure

In order to let participants reach their ceiling performance we introduced a 2-day setup for Experiment 2. The procedure of Experiment 2 was identical to that of Experiment 1 repeated over 2 days: during both days, participants completed a full 3-session 9-block experiment. Sequence of experimental condition blocks within a session was counterbalanced between sessions as well as between days.

#### 3.1.4 Data analysis

A three-way repeated measures ANOVA with a three-level factor Condition (Strong, Weak, Load-5), a three-level factor Session (1, 2, 3) and a two-level factor Day (1, 2) was used for statistical analysis. Greenhouse-Geisser correction was applied to the data which violated the assumption of sphericity. In the presence of significant interaction effects between factors, ANOVA was performed on each level of an interacting factor separately, with Bonferroni-Holm correction of p-values for multiple comparisons. Post-hoc pairwise comparisons were performed via paired-sample Student’s t-tests with Bonferroni-Holm correction for the noninteracting factors that reached significance.

### 3.2 Results

Experiment 2 aimed to investigate the dynamics of the training curve and its interaction with the effect of distractors on WM.

There was a statistically significant interaction between factors Condition, Session and Day on accuracy: F(4, 68) = 2.61, p < .04, eta_p_^2^ = .13 (Fig. 5). Therefore, two-way interaction and main effects of Condition and Session were analyzed for each day separately. Factors Condition and Session exhibited significant interaction in their effect on accuracy on Day 1 (F(4, 68) = 5.56, p < .01, eta_p_^2^ = .25) but not on Day 2. Given the significance of interaction, the main effect of Condition on accuracy during the first Day was analyzed for each level of Session separately.

**Figure 5.**
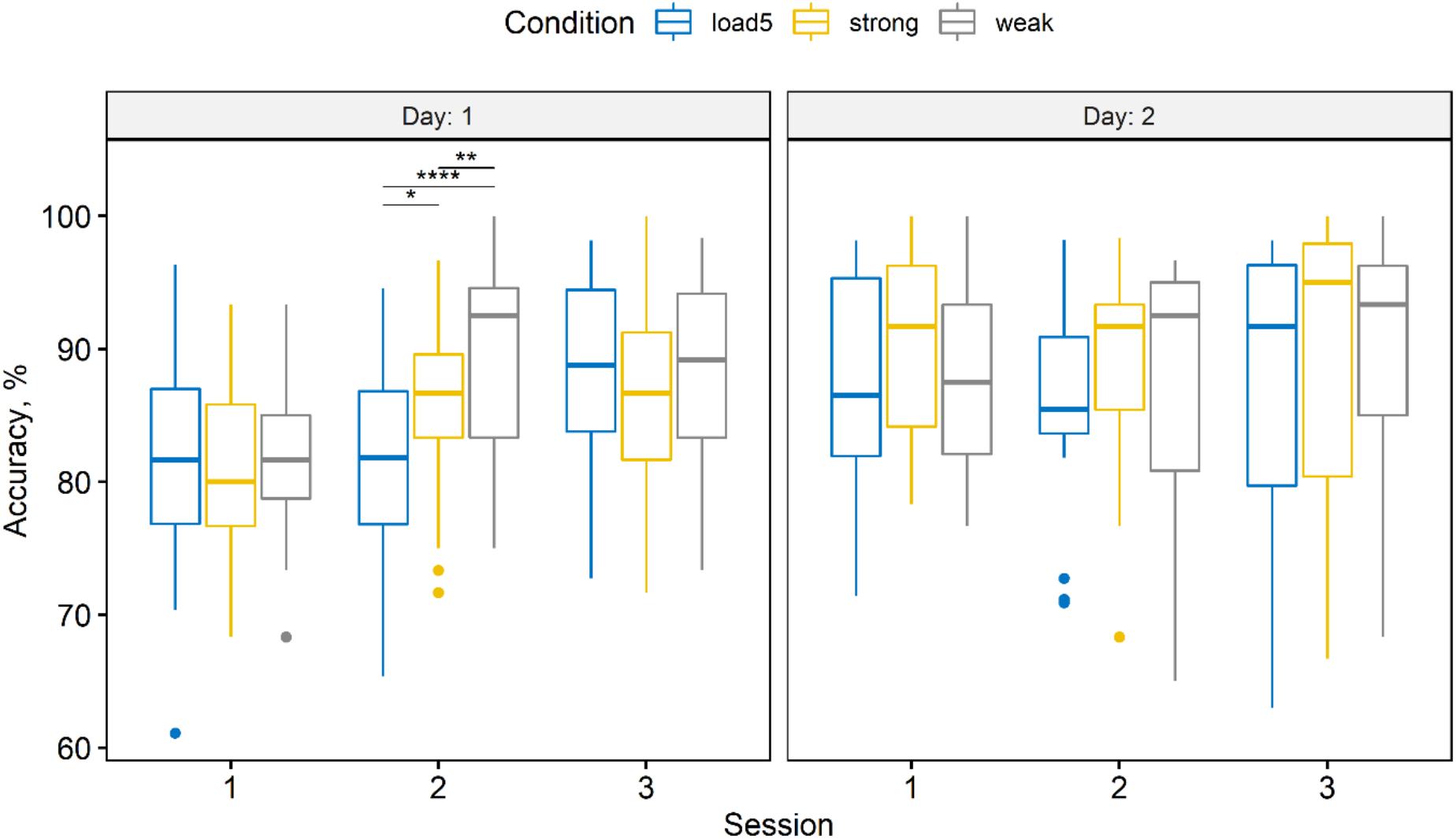
Experiment 2: Differences between distractor conditions, experimental sessions and days in accuracy. *p<.05,**p<.01,****p<.0001.

On Day 1, there was a statistically significant effect of Condition on accuracy during Session 2 only (F(2,34) = 16.9, p < .01, eta_p_^2^ = .5). Within that session, all three conditions showed significant differences (Strong-Weak: t = −3.3, p < .01; Strong-Load-5: t = −2.62, p = .02; Load-5-Weak: t = −6.03, p < .01). Factor Condition also showed a significant effect on accuracy on Day 2 (F(2,34) = 3.97, p = .03, eta_p_^2^ = .25). However, after correction of the p-value for multiple comparisons, the effect was not significant (p > .05).

Neither three-way interaction nor two-way interactions between factors Condition, Session and Day were significant in their effect on RTs. Therefore, we analyzed the main effects of the factors. RTs were significantly affected by factor Condition: F(2, 34) = 15.91, p < .001, eta_p_^2^ = .48 (Fig. 6). Upon pairwise comparisons, all three conditions showed significant differences (Strong-Weak: t = 4.43, p < .01, Strong-Load-5: t = −2.26, p = .02, Load-5-Weak: t = 7.19, p < .01).

**Figure 6.**
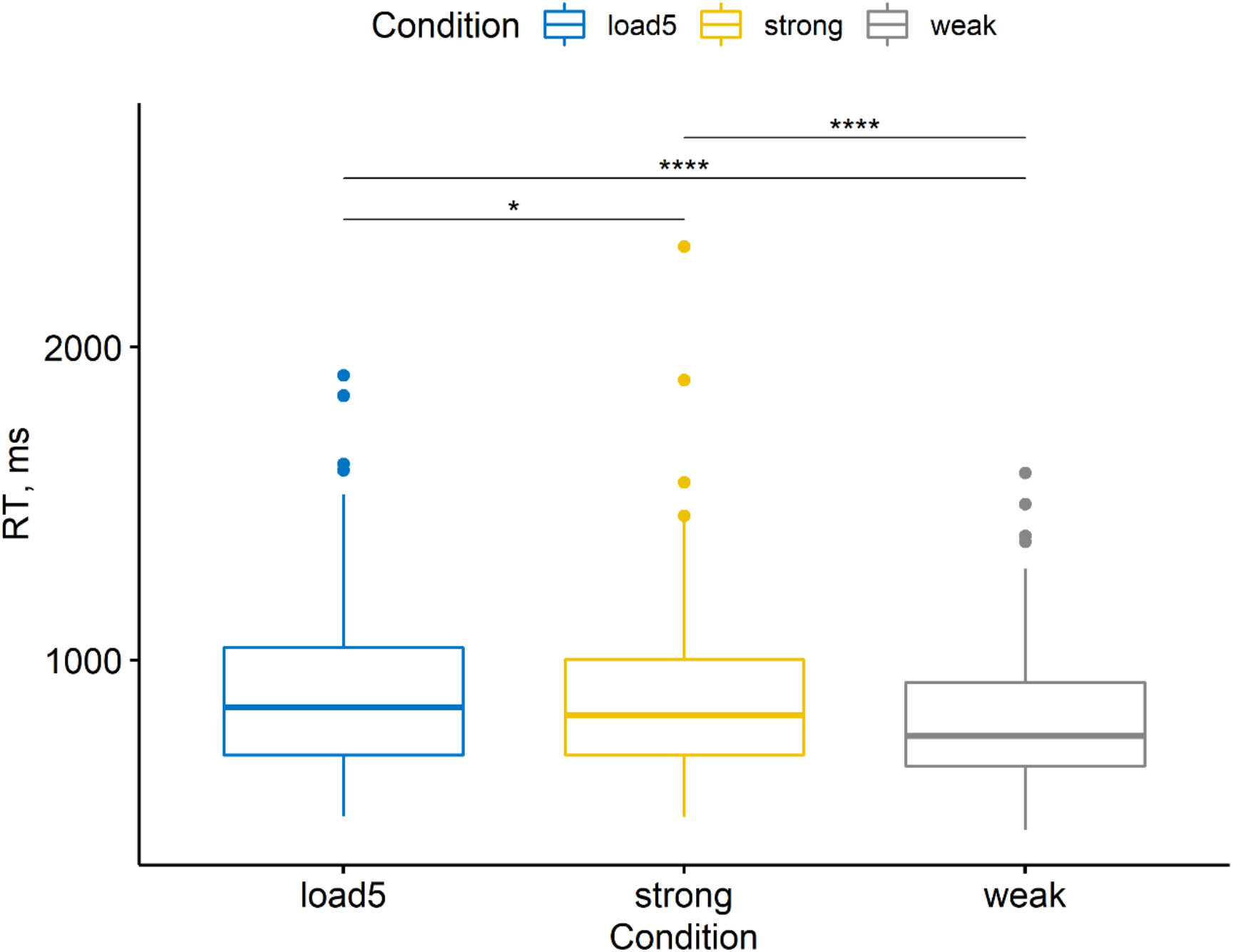
Experiment 2: Differences between experimental conditions in RTs. *p<.05,****p<.0001.

Additionally, we analyzed the effect of training on accuracy and RTs. Due to significant interactions between factors Session, Condition and Day on accuracy described above, we analyzed the effect of Session separately for each level of Day as well as for each level of Condition on Day 1. On Day 1, the effect of Session on accuracy was significant for each level of Condition (Strong: F(2,34) = 3.7, p = .04, eta_p_^2^ = .18; Weak F(2,34) = 12.5, p < .01, eta_p_^2^ = .42; Load-5: F(2,34) = 6.68, p < .01, eta_p_^2^ = .28). On Day 2, the effect of Session on accuracy was not significant. Session exhibited a significant effect on RTs irrespective of Condition or Day: F(2, 34) = 7.53, p = .002, eta_p_^2^ = .31 (Fig. 7).

**Figure 7.**
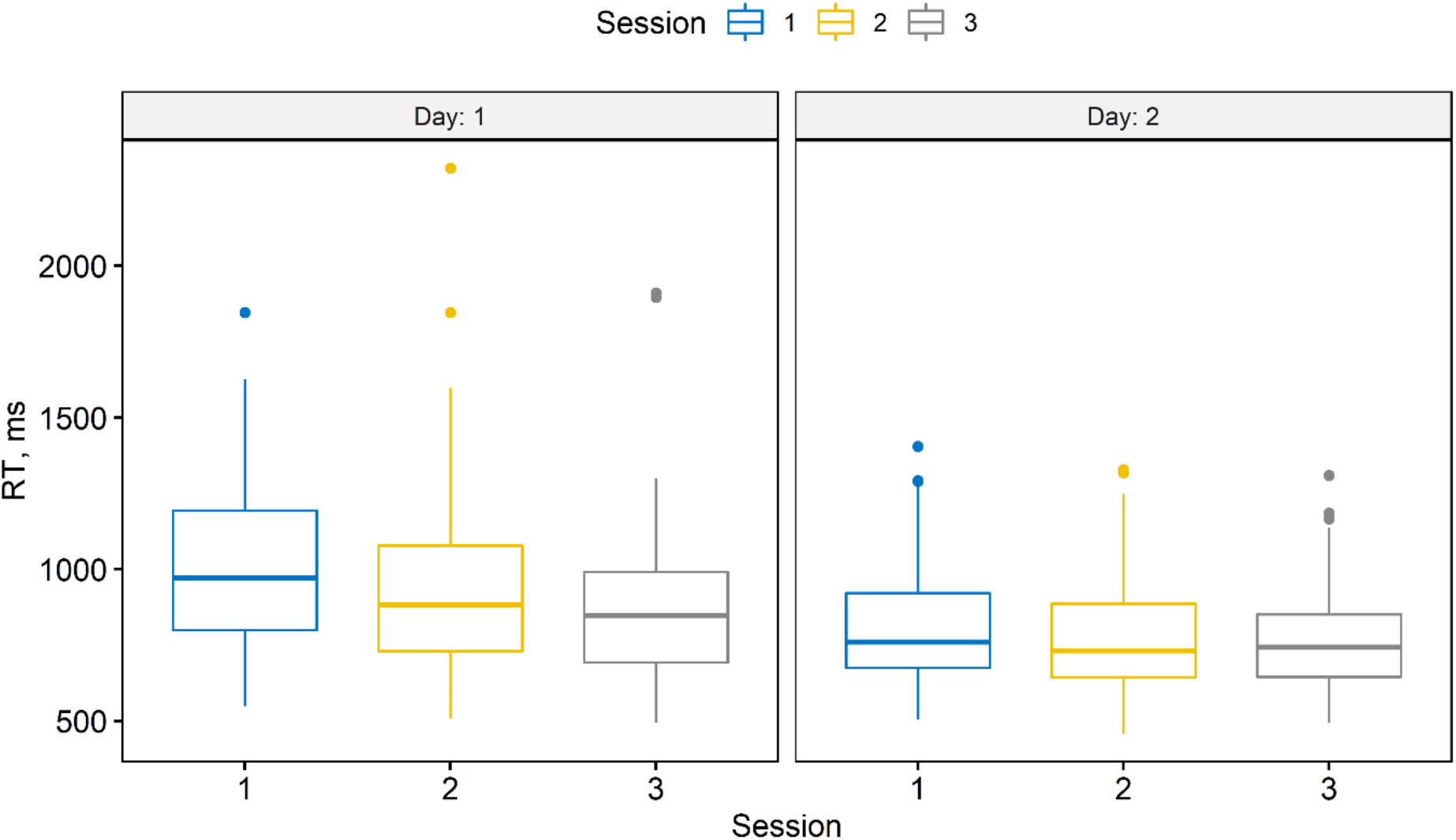
The difference in RTs between sessions and days.

### 3.3 Summary

Ceiling performance in accuracy was successfully reached by the second day of the experiment while RTs continued improving into the second day.

The effect of the distractor, as well as memory load, on accuracy appeared to be modulated by training: no significant difference between conditions could be detected after the point of peak performance was reached.

While RTs exhibited substantial improvement from training, it did not interfere with the effect of the distractor or memory load on RTs. The strong distractor condition resulted in longer RTs than the weak distractor condition while the condition with higher load and no distractor produced the longest RTs.

Overall, after performance across conditions reaches its ceiling performance, the effect of the distractor condition on accuracy appears to become marginal. The effect of conditions on RTs, on the other hand, appears to be robust towards training. Still, it is possible that 2-day training was not enough to reach ceiling performance in RTs.

## 4. Discussion

### 4.1 Novel stimuli in a WM task

Sternberg paradigm has been implemented in different WM domains with various types of stimuli. Bonnefond & Jensen (2012), who introduced a distractor to the Sternberg task, used letters of the English alphabet as stimuli. Usually, letters of a native alphabet are very well learned visual items for a participant. This familiarity is associated with distinct neuronal processes: stimuli that have prior representation in the brain are processed by a different neuronal network than novel ones (Xiang & Brown, 2004, Stern et al., 2001, Kirchhoff et al., 2000). Here, we introduced novel stimuli to the task - abstract figures, unfamiliar to participants and thus devoid of pre-established representations in memory. As a result, our stimuli were created such that the participants faced a task of memorizing, manipulating and ignoring items without engaging existing representations. This was likely reflected in a long learning curve of the present WM task.

### 4.2 Training effect

An important issue that arose during Experiment 1 was a substantial learning curve. Performance of the task significantly improved with time, independently from the main effects of experimental conditions (Fig. 4). Training effect has been previously documented in WM experiments (Pugin et al., 2015, Jaeggi et al., 2008, Schwarb et al., 2016). Potential problem lies in the possible difference of learning curves between experimental conditions. Experimental conditions of different complexity may be trained with different speed, which can in turn influence the observed difference between conditions, depending on when the conditions are compared. In Experiment 1, the training effect was present for most of the experiment duration but no interaction between distractor condition and duration of training was observed. Importantly, the results did not allow us to confirm that the ceiling stage of training has been reached during the experiment. It is possible that the ability to ignore a strong and a weak distractor, and the ability to memorize an additional item are trained differently. If so, behavioral success across different conditions may vary depending on the training stage.

### 4.3 Inhibition of novel figures during WM

The goal of Experiment 1 was to test the ability to retain and manipulate information in WM while ignoring external distractors irrelevant to the task. Importantly, the items for memorization and ignoring were unfamiliar to participants. Our findings showed the general effect of the distractor condition on both accuracy and RTs. However, upon pairwise comparisons, accuracy differed only between a higher memory load and a lower memory load with a weak distractor and not between different distractors or between a higher memory load and a lower memory load with a strong distractor. While these results are consistent with previous findings on the Sternberg paradigm with regard to different memory loads (Sternberg, 1996), they did not show any evidence of an effect of distractors on WM (Bonnefond & Jensen, 2012). RTs, on the other hand, revealed differences between different memory loads and between distractors of varying complexity but not between a higher memory load and a lower memory load with a strong distractor. These results may indicate that in the presence of novel stimuli, it takes as much effort to ignore a novel distractor as it does to memorize an additional item. Although the strong distractor condition employs a lower memory load, the need to ignore a distractor may require additional cognitive resources. Alternatively, participants may have failed to inhibit a strong distractor altogether, thus increasing memory load in an otherwise lower memory load condition. Both of these effects may be due to the novelty effect induced by unfamiliar stimuli.

### 4.4 Learning to inhibit

The goals of Experiment 2 were two-fold. Firstly, we aimed to investigate the effect of distractors on WM performance in the absence of a training effect, i.e. at peak performance.

Introduction of extended training period was sufficient to achieve ceiling effect in accuracy but not in RTs, which continued improving during the second day of experiment.

Accuracy did not reflect any differences between WM conditions after peak performance has been reached. Possibly, due to training, hindering effects of a strong distractor or a higher WM load became void. Previous findings indicated that training in visual filtering efficiency (ability to exclude irrelevant information from WM encoding) improves WM performance in individuals with low WM capacity (Li et al., 2017). After the training, the difference between conditions with and without distractors but with the same memory load was attenuated in comparison to pre-training results.

The second goal of the study was to observe possible differences between WM conditions as a function of training. Specifically, we focused on how training affects the inhibition process within WM. The effects of WM conditions on accuracy were significantly modulated by training. Significant differences between the conditions were present early on during the first day but diminished after ceiling performance has been achieved. As mentioned above, it is possible that more demanding conditions (strong distractor and higher WM load) benefited the most from training, while the easier condition did not change as much.

The effect of WM conditions on RTs, on the other hand, did not appear to be modulated by the training curve. RTs exhibited consistent dependency on experimental conditions across all sessions and days. While they improved with time, they did so equally for all conditions. Notably, while there was no significant difference in RTs between strong distractor and load-5 conditions during Experiment 1, it was present during Experiment 2 which included a smaller sample size but twice as many experimental sessions per subject.

Another explanation for the observed differences in performance between the two experimental days is memory consolidation. Memory consolidation is the process of transformation of shortterm memory contents into long-term representations, which happens over time and is facilitated by sleep (Dudai, Karni, Born, 2015). Thus, abstract figures, which were novel to participants during the first day of the experiment, may have been consolidated into long-term declarative memory by the second day. Relation of encoded stimuli to stored representations may relieve load on attention and WM (Oberauer, 2009). However, it may also disrupt filtering of distractors due to automatic encoding.

In conclusion, future studies targeting behavioral measures of WM performance as readouts of experimental intervention should address with care possible training effects that may mediate the results of interventions of different complexity. This issue is especially important with difficult tasks which may elicit longer training periods. In the present study, difficulty of the task was caused by introduction of novel stimuli. Moreover, d ifferent behavioral readouts (eg. accuracy and RTs, as in the present case) may behave differently with regard to this issue and may require different solutions. With respect to the present task, accuracy reflected differences between conditions better before the ceiling performance was achieved, while RTs were not as sensitive to training. Further investigation is needed to evaluate the effects of training on task performance in other commonly studied WM domains - visuo-spatial, auditory and other.

## Declaration of competing interests

The authors have no competing interests to declare.

## Acknowledgements

This work was supported by the Basic Science Program of the NRU Higher School of Economics. BG acknowledges partial support from CNRS, INSERM, ANR-17-EURE-0017, and ANR-10-IDEX-0001-02.

## Data availability

The data and materials for all experiments are available at request from the corresponding author.

## Supplementary materials

A. Power analysis

Power analysis was performed based on the results of Bonnefond & Jensen (2012).

B. Post-experimental questionnaire

Questionnaire at the end of the experiment (translation from Russian). Questions were asked and answers were noted down by the experimenter:

1. What memorization strategy did you use?
2. Did you change memorization strategy between blocks?
3. Did you change memorization strategy between conditions?
4. Did you have associations for symbols and distractors?

## Notes

### Competing Interest Statement

The authors have declared no competing interest.

